# Prolonged activity-deprivation causes pre- and postsynaptic compensatory plasticity at neocortical excitatory synapses

**DOI:** 10.1101/2023.07.05.547850

**Authors:** Derek L. Wise, Yasmin Escobedo-Lozoya, Vera Valakh, Berith Isaac, Emma Y. Gao, Samuel B. Greene, Aishwarya Bhonsle, Qian L. Lei, Xinyu Cheng, Stephen D. Van Hooser, Sacha B. Nelson

## Abstract

Homeostatic plasticity stabilizes firing rates of neurons, but the pressure to restore low activity rates can significantly alter synaptic and cellular properties. Most previous studies of homeostatic readjustment to complete activity silencing in rodent forebrain have examined changes after two days of deprivation, but it is known that longer periods of deprivation can produce adverse effects. To better understand the mechanisms underlying these effects and to address how presynaptic as well as postsynaptic compartments change during homeostatic plasticity, we subjected mouse cortical slice cultures to a more severe five-day deprivation paradigm. We developed and validated a computational framework to measure the number and morphology of presynaptic and postsynaptic compartments from super resolution light microscopy images of dense cortical tissue. Using these tools, combined with electrophysiological miniature excitatory postsynaptic current measurements, and synaptic imaging at the electron microscopy level, we assessed the functional and morphological results of prolonged deprivation. Excitatory synapses were strengthened both presynaptically and postsynaptically. Surprisingly, we also observed a decrement in the density of excitatory synapses, both as measured from colocalized staining of pre- and postsynaptic proteins in tissue, and from the number of dendritic spines. Overall, our results suggest that cortical networks deprived of activity progressively move towards a smaller population of stronger synapses.

**SIGNIFICANCE STATEMENT:** Blocking activity in neocortical slice cultures produced coordinated pre and postsynaptic changes at excitatory synapses. Functional and structural assays suggest that deprivation results in fewer excitatory synapses, but each is strengthened both pre- and postsynaptically. This may contribute to the emergence of epileptiform activity.

## INTRODUCTION

Homeostatic plasticity bidirectionally stabilizes firing rates of neurons in response to deviation from a set point (Turrigiano et al., 1998). Without this countervailing pressure, Hebbian synaptic plasticity would result in run-away hyperactivity or hypoactivity (Turrigiano and Nelson, 2004). Homeostatic plasticity is especially necessary as animals develop, to maintain properly balanced networks despite wide-ranging changes in synapse number and strength in early life (Bourgeois and Rakic, 1993; Cragg, 1975; De Felipe et al., 1997).

Despite the normally beneficial stabilizing influence of homeostatic plasticity mechanisms, there is also evidence that these same mechanisms can destabilize networks when engaged too strongly or for too long. Silencing activity *in vivo* with tetrodotoxin (TTX) infused into rat hippocampus (Galvan et al., 2000) and neocortex (Lee et al., 2008) results in lasting seizures. These maladaptive effects of activity deprivation only occur when begun early in life (e.g. during the second postnatal week) and when the deprivation is prolonged (typically 10-14 days). In these cases, compensatory plasticity can cause network excitability to “overshoot”, and this overshoot can persist. Homeostatic mechanisms offer a potential explanation for the frequent observation that seizures accompany developmental disorders, even in cases in which the disorders are known to initially cause reduced cortical activity (Nelson and Valakh, 2015).

To begin understanding the mechanisms that contribute to maladaptive plasticity in response to activity deprivation, we wished to understand in detail the changes that occur at excitatory synapses in mouse neocortical slice cultures at an age corresponding to the sensitive window for persistent seizures in response to activity deprivation *in vivo*. We studied an intermediate period of deprivation longer than 1-2 days, producing rapidly reversible homeostatic effects, but shorter than the roughly 10 days needed to produce lasting seizures. The main excitatory synaptic effect previously described for short-term (2D) deprivation is postsynaptic upscaling (Turrigiano et al., 1998), but additional mechanisms may be engaged during longer periods of deprivation. Presynaptic homeostatic plasticity has been observed in some studies, but not in others. This has been alternately attributed to multiple aspects of these studies. Possible explanations include: differences between hippocampal and cortical synapses (Mitra et al., 2011); differences between *in vivo* versus *in vitro* preparations (Echegoyen et al., 2007); or differences in the duration (Lindskog et al., 2010) or timing of deprivation (Wierenga et al., 2006). We examined this issue in the context of more extended periods of deprivation, because they can lead to lasting changes in circuit excitability and thereby to increased propensity for seizures (Valakh and Nelson, 2015). An additional potential source of discrepancy is that many prior studies have focused on dissociated culture preparations, in part because more intact preparations that preserve some or all features of tissue structure have been intractable to large-scale synapse analysis due to the high density of synapses and low optical and biochemical penetrability.

In this study we combined super resolution microscopy with custom analysis algorithms to analyze the density and size of excitatory synapses at scale in organotypic slice culture of mouse cortex following five days of silencing with TTX. The results revealed increases in the size of both presynaptic and postsynaptic compartments and these were confirmed in EM measurements. We also observed increases in the amplitude and frequency of miniature excitatory currents (mEPSCs). We also find a decreased density of excitatory synapses in our cultures, unexpected in light of the increase in mEPSC frequency. We conclude, therefore, that some excitatory synapses are lost, but those that remain are significantly strengthened pre- and postsynaptically. Therefore, synapses may compensate for the loss in release sites with enhanced release. These results suggest that prolonged deprivation of activity at neocortical excitatory synapses progresses over time to include changes to synaptic number and presynaptic strength, a departure from previous studies of two days or less of silencing which emphasized the role of postsynaptic regulation.

## METHODS

### Animals

All experiments were conducted in accordance with NIH guidelines for animal use and authorized by the Brandeis University Institutional Animal Care and Use Committee. C57BL/6 and CD1 strain mice were used for our experiments, with Ai14(RCL-tdT)-D Cre-conditional Td-Tomato mice (Jackson Laboratories Strain #007914) used for spine tracing and Emx1-Cre mice (Jackson Laboratories strain #005628) used for cell fill dendrite validation in our synapse imaging experiments. Our mice were fed *ad libitum* and housed at the Foster Animal Care Facility at Brandeis University.

### Organotypic Cortical Slice Culture

Brains were harvested from C57BL/6 mouse pups aged 7-9 days old and anesthetized with 40 μL of ketamine (20 mg/mL), xylazine (2.5 mg/mL), and acepromazine (0.5 mg/mL) mixture, administered intraperitoneally. Mouse forebrains were embedded in 2% low melting point agarose. Coronal brain sections were cut using a compresstome (Precisionary Instruments, Greenville, NC) in filtered ice-chilled artificial cerebrospinal fluid (ACSF; 126 mM NaCl, 25 mM NaHCO3, 3 mM KCl, 1 mM NaH2PO4 H20, 25 mM dextrose, 2 mM CaCl and 2 mM MgCl2, 315-319 mOsm) to 300 μm thickness and grown on Millipore Millicell inserts containing hydrophilic polytetrafluoroethylene (PTFE) membranes with 0.4 μm pores (#PICMORG50; Millipore Sigma, Burlington, Massachusetts) in 6-well dishes over media (1x MEM (Millipore-Sigma), 1x GLUTAMAX (Gibco; Thermo-Fisher Scientific, Waltham MA), 1 mM CaCl2, 2 mM MgSO4, 12.9 mM dextrose, 0.08% ascorbic acid, 18 mM NaHC03, 35 mM 1M HEPES (pH 7.5), 1 μg/mL insulin and 20% Horse Serum (heat-inactivated, Millipore-Sigma, Burlington, Massachusetts), pH 7.45 and 305 mOsm). A 1,000 units/mL PenStrep (Gibco; Thermo-Fisher Scientific, Waltham MA) & 50 μg/mL gentamicin (Millipore-Sigma, Burlington MA) antibiotic mixture was applied for the first 24 hours in a 35°C, 5% CO2 incubator. After changing into non-antibiotic media, media changes were performed every 2 days for the course of experiments.

For experiments relying on cell-wide expression of a fluorescent protein to track the morphology of dendritic spines, we used Ai14 mice with conditional expression of Td-Tomato. They then had a 1 μL drop of hSynapsin-dependent Cre (Addgene pENN.AA9V.hSyn.Cre.WPRE.hGH; #105553-AAV9) virus diluted 1:2000 from factory titer in ACSF deposited on the surface of the culture’s cortex to achieve sparse infection. The fluorescent marker was expressed at high levels in virus-infected cells.

For cultures undergoing developmental activity silencing, the sodium blocker tetrodotoxin (TTX, 500 nM; Tocris) was included in the media for five days, starting at equivalent postnatal day 12 (EP12) and lasting until EP17 (alternatively, DIV 5-10). This is a supramaximal concentration expected to stop action potential generation entirely.

### Electrophysiology

Cortical Slice cultures were carefully cut from the PTFE membranes and placed in a submerged slice chamber perfused with 33-35°C artificial cerebrospinal fluid (ACSF) containing 119 mM NaCl, 25 mM NaHCO3, 3 mM KCl, 1 mM NaH2PO4 H20, 25 mM dextrose, 2 mM CaCl and 2 mM MgCl2, adjusted to 315-319 mOsm. Layer 5 pyramidal cells were identified by their somatic morphology and targeted for whole-cell recording using an upright Olympus microscope equipped with a 40-X water-immersion objective. Their identity was corroborated in some cases by post-recording staining for biocytin contained in the internal solution. Whole-cell patch clamp recordings were obtained with glass pipette electrodes pulled from borosilicate capillaries (World Precision Instruments, Sarasota, FL) on a P97 Flame/Brown Puller (Sutter Instruments, Novato, CA). Pipettes had a tip resistance of 4-6MΩ and were filled with 100 mM K-gluconate, 20 mM KCl, 10 mM HEPES, 4 mM Mg-ATP, 0.3 mM Na-GTP, 10 mM Na-phosphocreatine. Miniature excitatory postsynaptic currents were measured at -70 mV in the presence of 0.1 μM TTX, 50 μM picrotoxin and 50 μM 2-amino-5-phosphonovaleric acid (APV). Only cells with R_in_ and V_rest_ maintained within 20% of their value ^5 min after break-in were kept. Currents were amplified using an Axoclamp 700B amplifier (Axon Instruments, Foster City, CA). Data were digitized at 20 kHz, filtered at 10 kHz, and collected using custom software written in IGOR Pro (Wavemetrics, Lake Oswego, OR) by Dr. Praveen Taneja.

### Electron Microscopy

Cultures in parallel conditions to previous experiments were used in accordance with methods described in (Studer et al., 2014) and (Bloss et al., 2016). A 2 mm disk was punched out from somatosensory cortex, and then the tissue was placed on HPF carriers and covered with incubation medium containing 20% dextran and 5% sucrose as cryoprotectant. Samples were high pressure frozen in a Leica HPM100 (Leica Microsystems, Vienna, Austria). This was followed by cryosubstitution in 96% acetone, 3% water and 1% anhydrous methanol containing 1% uranyl acetate (Electron Microscopy Sciences (EMS), Hatfield, PA), gradually raising the temperature from - 140°C to -45°C. Samples were embedded in Lowicryl HM20 Monostep resin (EMS) and polymerized under UV light for approximately 144 hours. Once embedded, 70 nm sections were cut and deposited on 200 mesh copper London Finder Grids coated with standard thickness formvar and carbon (EMS). The grids were counterstained with 4% Uranyl Acetate in 50% Ethanol and Lead Citrate (Venable and Coggeshall, 1965).

EM imaging and sample preparation were performed at the Brandeis Electron Microscopy Facility on a Tecnai F20 TEM (Thermo Fisher, Hillsboro, OR) operated at 200 kV and equipped with a Gatan Ultrascan 4kx4k CCD camera (Gatan, Pleasanton, CA). For large overviews of sections, we acquired montages of overlapping high-magnification images. Automated montage acquisition was facilitated by the microscope control software SerialEM (Mastronarde, 2005). Image processing, such as blending montages, was performed using the IMOD software package (Kremer et al., 1996).

### Synaptic Imaging

We performed an initial calibrating analysis on Emx-Cre mice (see Animals) expressing Cre-dependent GFP virus (Addgene pCAG-FLEX-EGFP-WPRE; #51502-AAV9) following a dot-wise application of 1:2000 diluted virus (see Organotypic Slice Culture methods). These mice subsequently expressed GFP sparsely in excitatory neurons in the cortex. Follow-up experiments were performed on C57BL/6 wild-type animals. Organotypic slices were fixed with 3% glyoxal for 30 minutes at 4°C and then 30 minutes at room temperature (Richter et al., 2018). Fixed slices were quenched in ammonium chloride/glycine solution (10 mM each) for 5 minutes, then washed 3 times into PBS and stored at 4°C until staining in PBS containing 1 μM sodium azide.

For staining, single-hemisphere slices were removed from PTFE membranes and first exposed to CUBIC 1 solution (Susaki et al., 2015) overnight at 37°C on a shaker. These slices were then treated with the DeepLabel Staining system (Logos Bio, Gyeonggi-do South Korea) suitable for thick samples, with overnight washes in each solution. Primary antibodies used to stain excitatory synapses were anti-PSD-95 (mouse, Synaptic Systems 124-011, Goettingen, Germany), VGLUT1 (guinea pig, Synaptic Systems 135 304), and anti-Bassoon (rabbit, Synaptic Systems 141 003).

Secondary antibodies were goat anti-mouse Alexa Fluor Plus 555 (Invitrogen A32727, Carlsbad CA), goat anti-guinea pig Alexa Fluor 647 (Invitrogen A21450), and goat anti-rabbit Alexa Fluor Plus 488 (Invitrogen A32732). All antibodies were diluted 1:2000 for use. In the final wash, DAPI (Thermo Fisher Scientific 62248, Waltham MA) was added at 1 μg/mL for 20 minutes at room temperature on a shaker to stain nuclei. Slices were carefully mounted with DeepLabel XClarity and allowed to set overnight before imaging.

Imaging of layer 5 of somatosensory cortex was performed on the Zeiss Airyscan confocal microscope system (Carl Zeiss, Jena, Germany). Raw 63x magnification images were pre-processed for Airyscan phase combination using Zen Black (Zeiss). Images were converted to single-channel TIFF files using FIJI (Schindelin et al., 2012). The subsequent analysis took place in a custom MATLAB suite developed by the Van Hooser lab and expanded for this purpose (Van Hooser, 2023).

The brightness of an object imaged through a scattering medium, such as brain tissue, decays exponentially with depth (**Supplementary Figure 2A**). To compensate for this, we measured the exponential decay, and then scaled by the inverse of this function. The best fit was found for a bi-exponential function. The signal was estimated from the intensities of the brightest 1% of pixels minus the noise estimated from the mean intensity of all pixels. The scaling factor applied was: DE_0_/DE_z_. Where DE_0_ is the signal intensity at the surface, DE_z_= a+b*e^-z/c^ + d*e^-z/f^, and z is the depth. Pixel intensities were shifted to avoid negative values introduced by noise subtraction. The dynamic range of our images was such that adequate contrast could be maintained after correction within the proximal 15 μm of the tissue, and we restricted our analysis to this depth.

We also took steps to compensate for the blurring introduced by the point spread function (PSF) of our microscope in X/Y through deconvolution (**Supplementary Figure 2B**). We calibrated this process using TetraSpeck fluorescent microspheres (Thermo Fisher Scientific T7279, Waltham MA) with 100 nm diameter (similar in size to a synaptic punctum) diluted 1:10,000 in water, dried, mounted with XClarity and coverslipped. Horizontal and vertical scans through images of each of 10 beads were averaged to compute a symmetric 2D kernel used to deconvolute the scaled images.

To identify puncta, we used thresholds reproducibly expressed as a percentile of the signal-to-noise of the depth-corrected image. To estimate the noise, the lower 75% of intensity values in each median filtered image were fit to a modified skewed Gaussian function: SG_i_ = a+b*e^-(x-c)2/(2*d2)^ * (1+erf((x-c)/sqrt(2)). (This differs from the skewed Gaussian distribution since b is not constrained to be 1/(2*sqrt(2*pi)*d)). The signal-to-noise ratio was the percentage of pixels of a given intensity (C_i_) above that expected based on the noise distribution: SN(i) = 100 * (C_i_-SG_i_)/C_i_.

Regions-of-interest (ROI) were identified by finding all pixels exceeding an upper threshold, and then a watershed algorithm was used to find connected pixels exceeding a lower threshold. Resegmentation allowed an ROI containing multiple peaks separated by lower values to be separated.

To optimize the upper and lower thresholds, we compared automated detection to detection of spines and puncta within spines by human observers who were blind to the experimental condition. Comparisons were performed on GFP-filled basal dendrites (see above, with Emx-Cre mice and CAG-flex-GFP virus) stained for presynaptic (VGLUT1) and postsynaptic markers (PSD-95). ROIs corresponding to dendritic spines were first selected manually, and then 1-3 human observers scored stained puncta (**Figure 3A**).

Using the human-selected regions-of-interest as a ground truth, we computed the probability of true positives and false positives for a variety of upper and lower thresholds. Values of 99.99 and 97 percentile maximized true positives and minimized false negatives in most of our datasets (**Figure 3B**). As a check, we examined the putative puncta occupancy rate of spines: no dataset exhibited a spine-puncta occupancy rate below 80% for either PSD-95 or VGLUT1, and most datasets had spine-puncta occupancy close to 100% for one or both channels (**Figure 3C**).

Our validated threshold was then used on the full data set comprising X/Y dimensions of 72 × 72 μm, in a 3 μm section with 0.2 μm increments (16 z-frames in total) selected from within the most superficial 15 μm in each slice, for a total volume surveyed of 15.560 cubic microns. Four nearby area replicates were surveyed for each sample. To avoid apparent noise artifacts which typically involved puncta with one or more linear dimensions occupying only a single pixel, a size filter was set to 2^3, or 8 cubic pixels. Adjacent but distinct puncta were separated with a watershed algorithm. Pre- and postsynaptic puncta were considered colocalized if they had overlap >1% of their total volume with an X-Y jitter of 2 pixels in each direction, and no jitter in the Z-dimension (since Z resolution is much less precise in confocal systems than it is in X-Y). Colocalized puncta were quantified in terms of their 2D cross sectional area in the plane containing their brightest pixel, as well as by overall volume (not shown).

### Cell Count in Organotypic Cultures

Cells were counted at 10x from the same regions of the same sections used for synaptic imaging, also on the Zeiss Airyscan system. A 461 × 461 μm area was imaged through the full thickness of the slice in 2.5 μm increments for each sample, and DAPI-positive nuclei were counted in CellProfiler (Stirling et al., 2021) using an adaptive Otsu threshold (correction factor 0.9 - 1.0, expected diameter 20-60 pixels). Each nucleus was only counted once, in the plane in which it was best in focus. The total nuclei counted in the sampling area were normalized by the thickness of the slice to obtain a nuclei density for each slice. Slice thickness was measured post-fixation and post-histology in mounted slices, by imaging through each slice until the last layer of nuclei that was in focus was imaged. Dehydration due to sample processing led to typical mounted slice thicknesses ranging from 15 to 35 μM; living slices are estimated to be ^100 to 150 μM based on their thickness measured after cryofixation, which does not lead to any sample dehydration.

In a subset of our slices, synaptic puncta and nuclei were imaged and counted in the same region to obtain the nuclei normalized synaptic density. Briefly, this normalization was done by dividing the total number of colocalized pre- and postsynaptic puncta (putative synapses) found in the optimal imaging region by the number of μm^3^ in the imaged volume (measured with the calibrated Airyscan), and then dividing this synaptic density by the average nuclei density found within the same slice.

### Statistical Analysis

For comparisons between two conditions, two-tailed student’s t-tests were used when the distributions were not highly skewed (skewness between 1 and -1). If the variance between groups was significantly different, a Welch’s correction was used. For comparisons in which one or both distributions had a greater magnitude skewness, a Mann-Whitney (also called Wilcoxon Rank Sum) test was used.

## RESULTS

### Prolonged activity deprivation causes an increase in mEPSC amplitude and frequency

To study homeostatic plasticity at excitatory synapses in response to activity deprivation we prepared organotypic slice cultures that included neocortex at P7 and cultured them for 10 days. During this period the neurons reform connections and spontaneous bouts of activity, termed up states, emerge (Johnson and Buonomano, 2007). To broadly block activity, we applied the voltage-gated sodium channel blocker TTX (0.5 μM) to a set of cultures throughout the last 5 days of this 10-day period and compared properties of excitatory synapses to age-matched controls with uninhibited spontaneous activity. This manipulation induces a robust homeostatic program that profoundly shifts network dynamics towards hyperexcitability (Koch et al., 2010; Wise et al. 2023). We obtained whole cell patch clamp recordings from layer 5 pyramidal neurons in somatosensory cortex. Analysis of pharmacologically-isolated AMPA-driven mEPSCs revealed that the 5-day activity deprivation caused a 47% increase in the amplitude of spontaneous excitatory currents (**Figure 1**, CTRL 11.7±5.4 pA, n = 15, TTX 17.2±5.0 pA, n = 25 cells) which was highly significant (Mann-Whitney p = 0.002), consistent with previous reports of synaptic scaling (O’Brien et al., 1998; Turrigiano et al., 1998). However, unlike in most prior work performed in dissociated cortical cultures after two days of deprivation, we observed a dramatic 2.2-fold increase in mEPSC frequency (CTRL 2.5±2.5 Hz, n = 15, TTX 5.6±2.8 Hz, n = 25 cells) which was also highly significant (Mann-Whitney p < 0.001). This suggests that after 5 days of deprivation, both the pre- and postsynaptic strength of excitatory neocortical synapses are increased.

**Figure 1.**
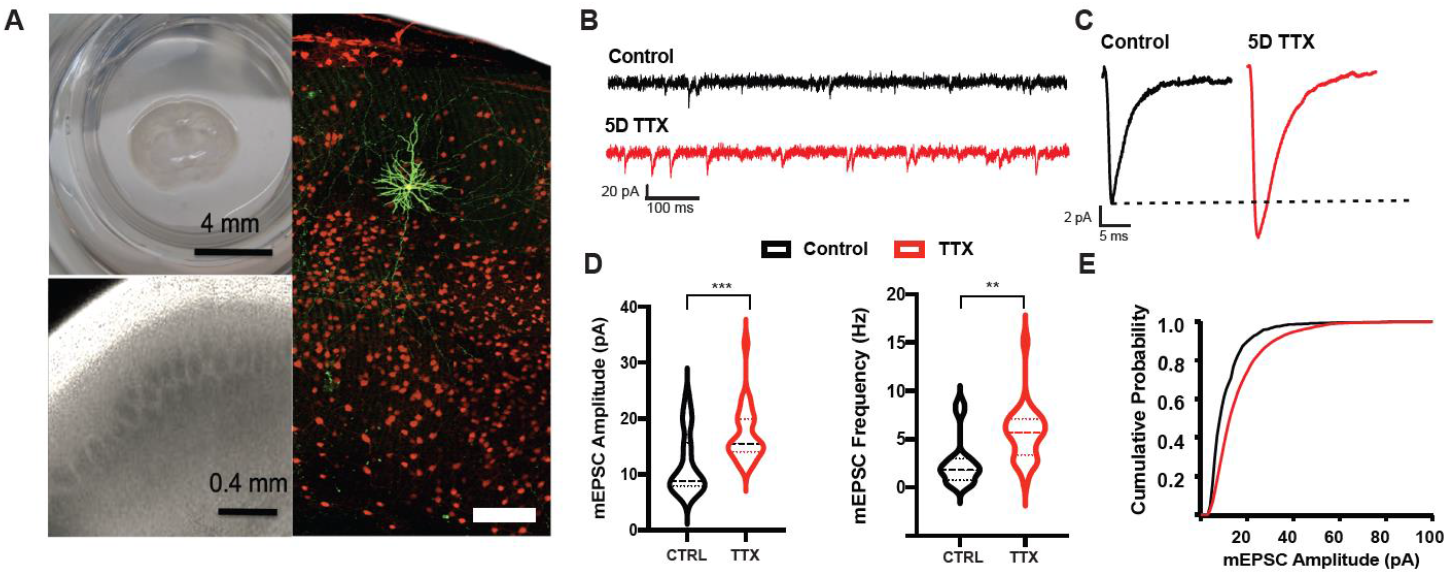
Spontaneous synaptic currents increased in amplitude and frequency following five days of activity deprivation. **A**. Coronal slice culture including cortex (left, top); 4x brightfield image showing cortical layers (left, bottom); fluorescent image of culture with filled pyramidal neuron (green) at P17/DIV 10 (right). **B**. Example miniature EPSP traces of age-matched control whole cell voltage clamp and 5D TTX recordings, one second each. **C**. Average mEPSCs, compiled from the average trace of 100 minis from each cell for deprived and control samples. **D**. Average amplitude (left) and frequency (right) of mEPSCs for each cell (n = 15 CTRL, 25 TTX cells from 7 and 8 independent slice cultures). **E**. Cumulative histogram of amplitude for subsampled minis used to compile the average mini in **C**; TTX-treatment shifted the curve to the right of control.

Quantitative assessment revealed other changes in cellular and synaptic properties. There was no change in measured whole cell capacitance (53.5±14.9 pF CTRL, 40.9±8.9 pF, p = 0.40) but input resistance at rest was increased (297.3±23.5 MΩ CTRL, 580.3±96.0 MΩ TTX, p = 0.012) consistent with other observations of increased intrinsic excitability of activity-deprived neurons (Desai et al., 1999; Sokolova and Mody, 2008). Although the rising phase of the recorded minis did not differ by condition (10%-90% rise time = 3.9±0.06 ms CTRL, 4.0±0.09 TTX, p = 0.38); the decay kinetics were prolonged (decay ***τ*** = 4.8±2.0 CTRL, 3.7±1.9 TTX, p = 0.005) perhaps reflecting a change in the auxiliary or receptor subunit composition with prolonged deprivation.

### Length of post-synaptic density is increased following activity deprivation

Although most prior work on neocortical homeostatic plasticity has emphasized the primarily postsynaptic change (for review see Turrigiano and Nelson, 2004), several prior studies on homeostatic responses in hippocampal dissociated culture (Murthy et al., 2001), slice culture (Mitra et al., 2011) and in vivo (Echegoyen et al., 2007) have emphasized an additional or greater component of this plasticity which is presynaptic (Delvendahl and Müller, 2019). In some cases, this increased release has been shown to correlate with increases in the size of the presynaptic terminal and in the number of docked vesicles (Murthy et al., 2001).

To describe the morphological correlates of the observed homeostatic plasticity in neocortical slice culture, we visualized synapses in untreated cultures or cultures treated with 5D of TTX using electron microscopy after high-pressure freezing and freeze-substitution. Cryofixation achieves the most native-like preservation of brain tissue, avoiding fixation artifacts (Kellenberger et al., 1992), but also leads to lower contrast and difficulty with visualization of postsynaptic densities at all excitatory synapses. To ensure measurements include only excitatory synapses, we restricted our analysis to synapses made onto dendritic spine heads (Bartol et al., 2015). All measurements were made blind to condition.

We first measured the size of the postsynaptic compartment and observed a dramatic shift in the distribution of synapse lengths at postsynaptic densities of activity deprived cultures (Fig. 2B; from 260.0±14.0 to 468.4±29.2; Wilcoxon signed rank test p < 0.001). This increase is expected, given the increase in mEPSC amplitude observed, but the mean increase of 80% was larger than the 47% increase observed physiologically.

**Figure 2.**
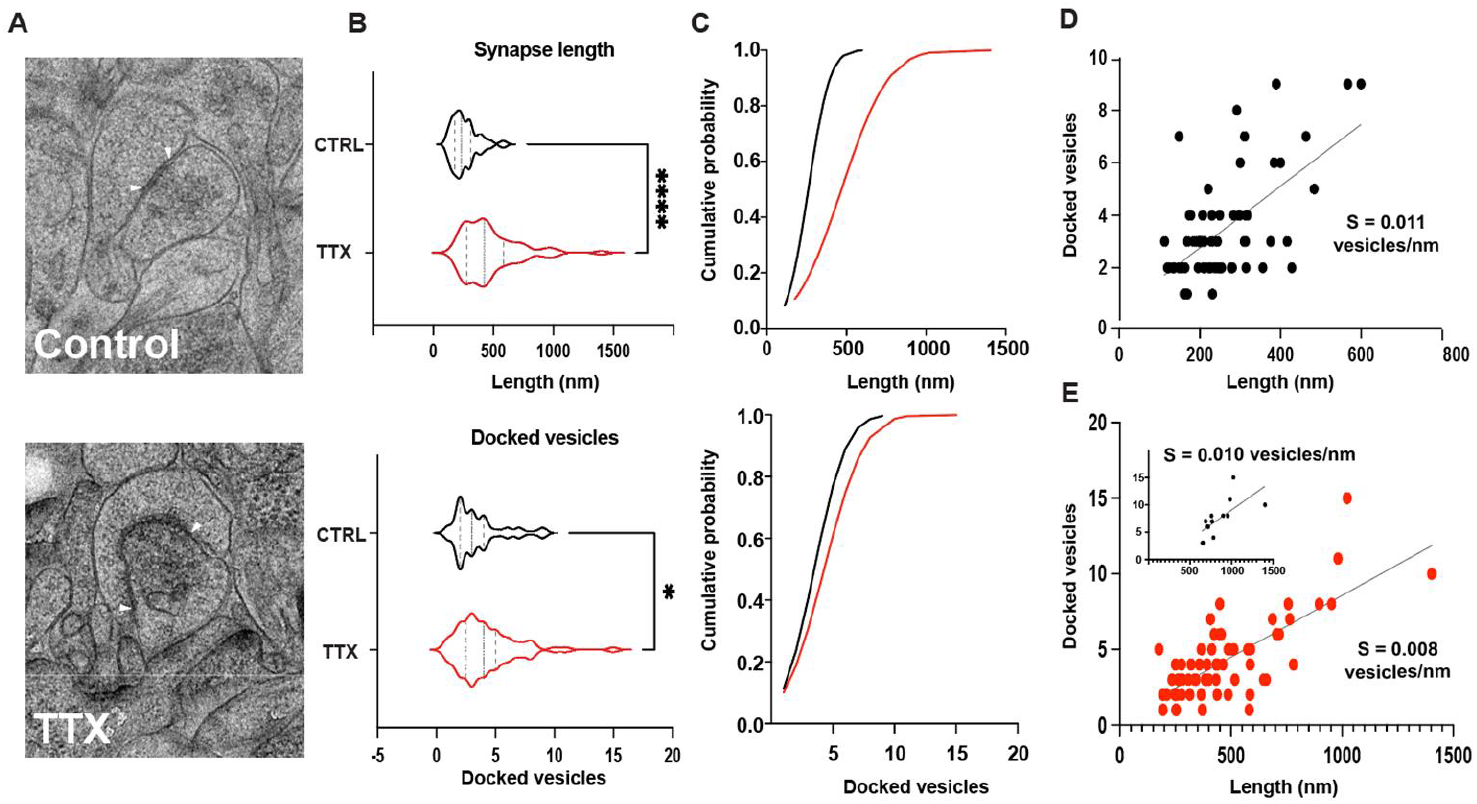
Electron microscopy images show enlarged synapses with more docked vesicles. **A**. Example electron micrographs from control and slices silenced for 5 days, with the edges of synapses denoted with white arrows; the circumference between these arrows is the synapse length. Scale bar is 500 nm. **B**. Counts for synapse length, and count of docked vesicles at these synapses (n = 62 synapses from 8 CTRL cultures, 67 synapses from 7 TTX-treated cultures, Wilcoxon signed rank test p < 0.001). **C. (**Top) Cumulative distributions of PSD length for synapses in B. (Bottom) Cumulative distributions of docked vesicles at synapses in B. **D & E**. Relationship between length of the postsynaptic density and docked vesicle counts for CTRL (D) and 5D TTX (E).The straight lines are linear fits with variable intercept, the slope of the relationship (S) decreases after 5D in TTX. **Inset**. As **E**, but separated but only for the longest synapses (> 600 nm). Most of the change in slope derives from measurements in the smallest synapses as the slope for the longest synapses (inset), right (s = 0.010 vesicles/nm) is nearly identical to the control slope (s = 0.011 vesicles/nm).

**Figure 3.**
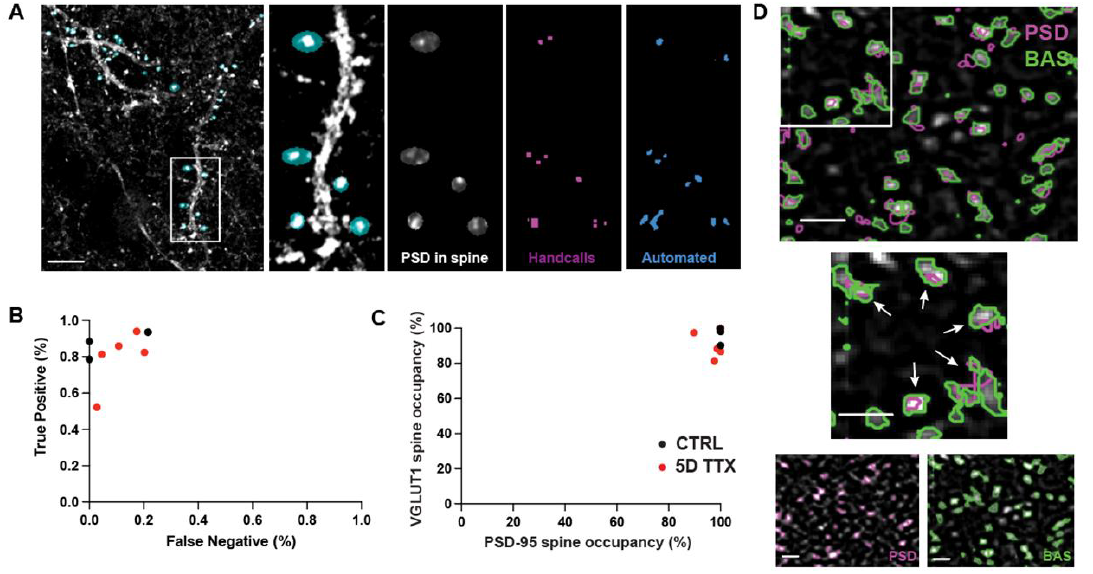
Calibration of synapse identification based on axospinous contacts. **A**. Sample cell fill containing dendritic branches (white) with spine ROIs selected manually (cyan ovals). One area of the left-hand image is emphasized on four right-hand panels, where spine ROIs contain PSD-95 signal, with hand-called puncta and automatic threshold ROIs. **B**. Optimal thresholding algorithm (99.99%) as compared to human-made selections of the same channel within labeled spines (black = CTRL and red = 5D TTX), with true positive (puncta caught by both the experimenter and automated threshold) and false negative (puncta called by the automated threshold but not the experimenter) rates. **C**. Quantification of PSD-95 and VGLUT1 puncta occupancy in dendritic spine heads for CTRL (black) and TTX (red) samples. **D**. Example colocalization of presynaptic Bassoon (green) and postsynaptic PSD-95 (magenta), with individual channels (left two images) and the union of the two (right two images; zoomed in on rightmost). White arrows highlight the location of colocalized presumptive synapse locations. No somas are present in this example. Scale bars all represent 1 μm.

In order to quantify presynaptic structure, we measured the number of docked vesicles at the portion of the presynaptic terminal apposed to the PSD. We found that in TTX-treated slices, the number of docked vehicles increased by 21% (from 3.48±0.27, n = 60 synapses from 8 CTRL cultures, to 4.23±0.32, n = 65 synapses from 7 TTX treated cultures, Wilcoxon signed rank test p < 0.001). This observation is consistent with prior studies in hippocampus (Murthy et al., 2001) and with our observation of increased mEPSC frequency suggesting that structural changes arising from 5 days of full activity deprivation occur at both sides of the synapse to comparable degrees.

### Activity deprivation increases volume and intensity of the presynaptic and postsynaptic compartments

Although electron microscopy with cryofixation provides a “gold standard” measurement of synaptic morphology (Studer et al., 2014), our implementation was necessarily relatively low throughput and allowed us to measure changes at only a modest sample of cortical synapses. To provide a complementary view permitting lower resolution but more complete sampling of cortical synapses, we turned to super-resolution light microscopy. Using the same 5-day TTX treatment, we immunostained the presynaptic compartment with pan-synaptic Bassoon and excitatory-specific VGLUT1 while also highlighting the postsynaptic compartment with excitatory-specific PSD-95. Resolving individual synapses in highly dense neocortical tissue was aided by super resolution imaging with an Airyscan microscope. This technique allowed us to visualize endogenous proteins in dense areas without the drawbacks associated with overexpression of tagged proteins (Sauerbeck et al., 2020; Wu and Hammer, 2021).

Light microscopic analysis comes with reduced certainty that visualized synaptic proteins are located exclusively at synapses. In order to calibrate our method, we chose first to examine a population of presumptive synapses localized to dendritic spines visualized in GFP-labeled neurons (**Figure 3**). We presume that the vast majority of spines contain at least one synapse (Arellano et al., 2007) and that most cases of colocalized pre- and postsynaptic proteins, in which the postsynaptic protein is located within the spine head, correspond to synapses. Because detection of synaptic puncta depends critically on setting thresholds to distinguish them from background, we sought a reproducible method for setting thresholds based on a fit to the underlying noise distribution. Three human observers selected spine heads while blinded to condition, as well as the PSD-95 and VGLUT1 signal within the spine head area (**Figure 3A**). These hand-called synaptic puncta were then used to choose thresholds (see Methods; **Supplemental Figure 1B**) that maximized true positives (called by human observers and the automated threshold) while having relatively low levels of false negatives (puncta called by the automated threshold but not by the observers; **Figure 3B**). These automated thresholds also resulted in high observed spine head occupancy for both PSD-95 and VGLUT1 puncta (80-100% for both compartments, **Figure 3C**).

Using this validated automated threshold procedure, we then examined a larger number of samples that did not contain cell fills or hand-called spines (**Figure 3D**). Instead of using VGLUT1 for the presynaptic label, we used Bassoon for this larger analysis as it was a more consistent label in our hands. These images were used for our quantitative analysis, allowing for quantification of tens of thousands of putative synapses per condition (**Figure 4**). Presumptive synapse size, measured as the 2D area in the brightest frame of each pair of colocalized puncta, was calculated for both synaptic compartments, showing a 21% increase in area for PSD-95 (**Figure 4D**; from 99.6±4.8 to 121.0±6.0 nm^2^, Two-tailed T-test, p = 0.018) and a 14% increase for Bassoon (from 128.0±3.7 to 145.5±6.4 nm^2^, p = 0.038), consistent with the increased function measured physiologically and the increased synaptic size measured ultrastructurally. Increases in mEPSC frequency, as observed in **Figure 1**, could potentially reflect a combination of an increased number of synapses and increased release at each synapse. Surprisingly, the density of PSD-95 puncta colocalized with Bassoon puncta decreased by 34% over 5D of TTX treatment (**Figure 4B**; p = 0.046; from mean 0.83±0.10 to 0.55±0.06 colocalizations per μm^3^). This suggests that, in this preparation, prolonged activity deprivation reduces the overall number of excitatory synapses, but makes each stronger enough to functionally outweigh this reduction.

**Figure 4.**
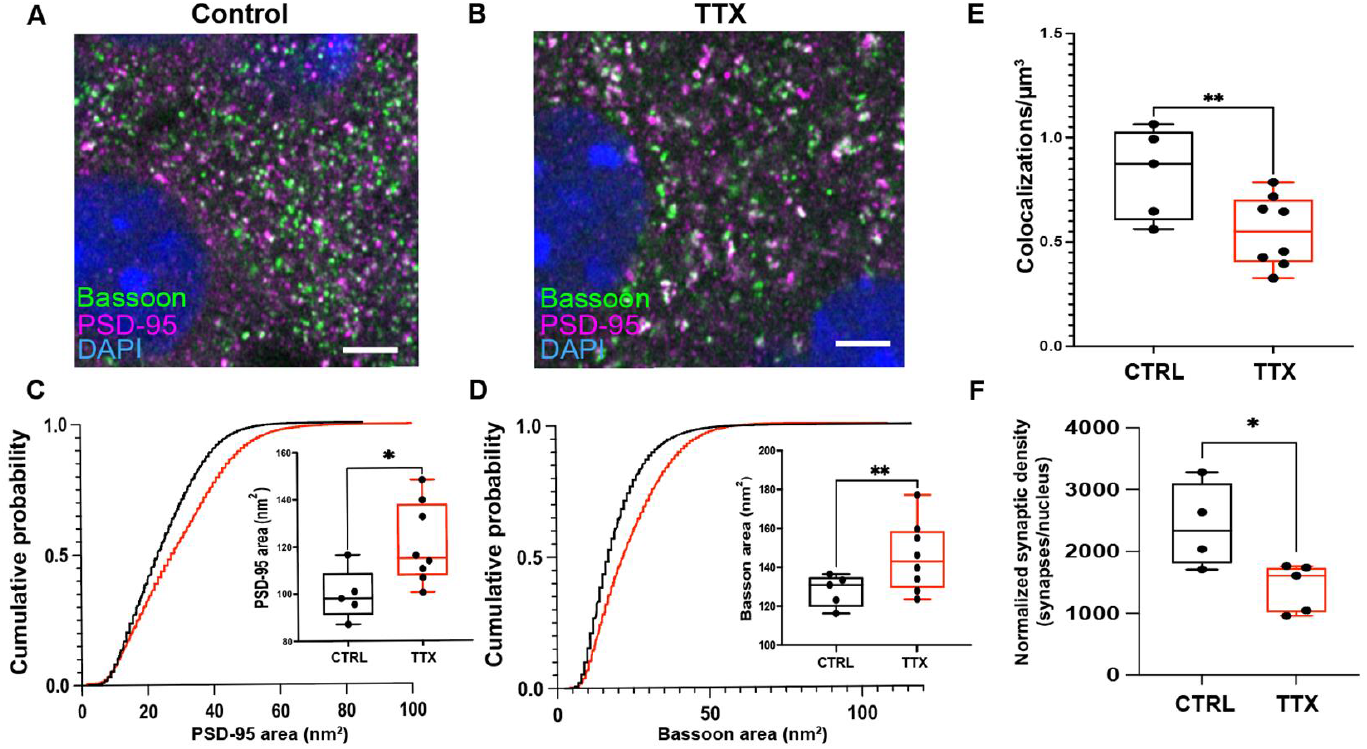
The size of excitatory synaptic compartments increased with silencing, while the density of excitatory synapses decreased. **A, B**. Representative images of L5 somatosensory cortex near the slice culture surface taken on Zeiss Airyscan microscope at 63x, with presynaptic Bassoon (green), postsynaptic PSD-95 (magenta), and nuclear stain DAPI (blue). Scale bar 100 micrometers. Colocalization in inset, top left, at white arrows. Scale bar 5 micrometers. **C**. Average empirical cumulative distribution of 2D cross sectional area of postsynaptic PSD 5 CTRL and 8 TTX samples, with inset showing sample means (stars denote 2-tailed T-test p < 0.05; double stars denote p < 0.01). **D**. As C, for Bassoon. **E**. Decreased synapse density (n = 5 CTRL, 8 TTX slices) of PSD-95 puncta per cubic micron colocalized with Bassoon following prolonged silencing (CTRL in black, TTX in red). Stepwise rise reflects that only discrete integers are possible for cross sectional area (in pixels). **F**. Synapse density normalized by the cell density in the same region of the slice where synapses were measured is reduced after TTX (n = 4 CTRL, 5 TTX slices).

The decrease in synapse density and increases in pre- and postsynaptic size we observed were also born out in our spine-validated calibration dataset that used VGLUT1 instead of Bassoon as a presynaptic marker, albeit with an underpowered sample size (n = 3 CTRL and 5 5D TTX stacks from 2 animals). This smaller sample showed a 34% decrease in PSD/VGLUT1 colocalization density (p = 0.093), a 41% increase in colocalized PSD-95 2D area (p = 0.024), and a 19% increase in VGLUT1 2D area (p = 0.11). These findings corroborate our somewhat surprising results with a slightly different readout.

The loss of colocalized synaptic proteins was also comparable to the changes in all of the PSD-95 and Bassoon puncta, including the ones that were not colocalized with an appropriate synaptic partner. Here too, we found decreased density with TTX - a 21% decrease in PSD-95 density (p = 0.198) and a 37% decrease in Bassoon density (p = 0.009), as compared to the 34% decrease in putative synapses from colocalizations seen above. In this broader population, a 21% increase (the same as found for colocalized puncta) was apparent for PSD-95 area (p = 0.049), with a 9% increase in Bassoon area (p = 0.070) as compared to a 14% increase from colocalized puncta above.

### Overall cell density does not change with application of tetrodotoxin

Synaptic density decreases could reflect cell death during the period of deprivation. To determine if the decrease in synaptic density reflects a decrease in the density of neurons, we examined DAPI-stained volumes at lower resolution over a much larger region (10x magnification; 461 × 461 μm area over the full Z-extent of each slice) in the same layer 5 region of somatosensory cortex from which our synapse count data were obtained (Figure 4). Nuclei were classified and counted using Cell Profiler software (Stirling et al., 2021). The resultant nucleus density, which includes non-neuronal cell types such as glia, was compared across conditions. We found no changes in total nuclei or slice thickness between age-matched control and TTX-treated samples (total nuclei CTRL 1910 ±103, n=6; TTX 2249± 63, n= 6 slice pairs, Two-tailed T-test p = 0.064; **Supplemental Figure 2B & D**) suggesting the decrease in synapses in TTX-treated slices does not result from cell loss. We observed a non-significant trend for TTX-treated slices to be thinner than control slices (slice thickness: Control 24.8 ±1.8, TTX 22.1 ± 0.4; n=6 slice pairs; t-test p = 0.28, Supplemental Figure 2B). The average number of nuclei in each Z-frame did show a slight increase with added TTX (average nuclei: CTRL 397.4±16.6, TTX 460.0 ± 6.1; n = 6 slice pairs; Two-tailed T-test p = 0.005, **Supplemental Figure 2C**), but this was in the opposite direction of what we would expect if this effect were to account for the observed loss in synapse density. To more directly address the relationship between nuclear and synaptic density, we plotted these together (**Supplemental Figure 2E**) and found only a weak relationship (r^2^ = 0.16). Synaptic density did not appear to reflect a lower cell density, since normalizing the synapse density to the density of nuclei, where both were measured, still exhibited a significant decrease with the application of TTX for five days (CTRL mean 2416 synapses/nucleus, TTX men 1425 synapses/nucleus; Two-tailed T-test p = 0.029).

### Activity deprivation shows a decrease in spine density

Since we observed an unexpected reduction in synapse density in our light microscopic synaptic imaging, we wondered if deprived neurons had fewer synaptic spines. In TdTomato-filled layer 5 pyramidal neurons, we measured the density of basal dendritic spines per unit length of dendrite (**Figure 5A**), finding a significant 14.2% decrease (**Figure 5B**, from 0.72±0.02 to 0.62±0.02; n = 18 CTRL, 19 TTX dendritic segments (each from a different neuron) from 4 animals, Two-tailed T-test p = 0.024). Most cortical excitatory synapses are thought to occur on spines (Bartol et al., 2015). Therefore, a reduced number of spines is consistent with an overall reduction in excitatory synaptic density following prolonged silencing.

**Figure 5.**
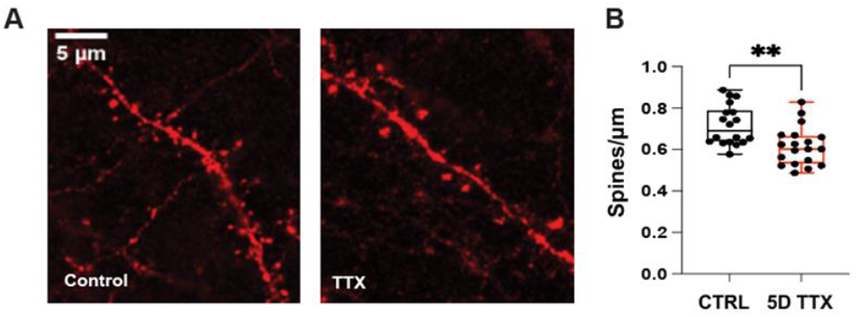
Lower spine density in basal dendrites from silenced cultures. **A**. Neurons in slice cultures from Ai14 mice carrying a Cre-dependent TdTomato allele were sparsely labeled by lentiviral Cre delivery. Representative image (left) shows a dendritic segment from a basal dendrite of a tdTomato-labeled neuron in layer 5 of a control slice. Representative image (right) shows a basal dendritic segment from a neuron in Layer 5 of a slice treated with TTX for 5 days. Dendrites run from top left to bottom right; other fluorescent signals arise from nearby axons. **B**. Basal Dendrite Spine quantification (n = 18 CTRL and 19 TTX dendritic segments, p = 0.0024).

## DISCUSSION

Excitatory synaptic strength is homeostatically regulated in response to deprivation of ongoing activity (Turrigiano and Nelson, 2004). Initial studies at cortical synapses in dissociated cultures emphasized that this homeostatic plasticity at excitatory synapses initiates rapidly (Ibata et al., 2008), is readily reversible (Koch et al., 2010), and is expressed primarily or exclusively as a postsynaptic change in the abundance of glutamate receptors and associated proteins (Ehlers, 2000; Gainey et al., 2009; Turrigiano et al., 1998). Although this “synaptic scaling” occurs at a variety of central synapses, such as those in spinal cord (O’Brien et al., 1998) and hippocampus (Lissin et al., 1998; Shepherd et al., 2006; Stellwagen and Malenka, 2006), several studies at hippocampal synapses have suggested that these postsynaptic changes can also be accompanied by, or are eclipsed by, presynaptic changes in excitatory synaptic transmission. These include studies of dissociated cultures (Burrone et al., 2002; Murthy et al., 2001; Tokuoka and Goda, 2008), slice cultures (Mitra et al., 2011) and acute slices following activity blockade in vivo (Echegoyen et al., 2007). Here we have reexamined this issue at neocortical synapses in slice culture and conclude that with a longer period of deprivation, of five days instead of two, presynaptic changes are quite prominent. Specifically, we found both physiological evidence (increased mEPSC frequency) and electron- and light-microscopic morphological evidence consistent with enhanced presynaptic function and size (increased synapse length, increased docked vesicles and increases in the size of puncta stained for presynaptic proteins). Taken together, our findings are consistent with the idea that activity deprivation homeostatically upregulates excitatory synapses both pre- and post-synaptically. The discrepancy with prior reports of a purely postsynaptic change could reflect a difference between dissociated culture and slice culture, or could reflect the longer period of deprivation. The former is more likely, since a recent study in slice culture also reported evidence for pre- and postsynaptic changes after two days of activity deprivation (Valakh et al., 2021).

Surprisingly, we found that this enhanced presynaptic function occurred in the setting of an overall loss of excitatory synapses. Although our choice of cryofixation with freeze-substitution did not permit us to verify reduced synapse density ultrastructurally, two light microscopic methods supported this conclusion. First, we identified putative synapses by colocalizing presynaptic Bassoon or VGLUT1 staining with postsynaptic staining for PSD-95. This method confirmed the fact that collections of synaptic proteins, like the synapses themselves identified in the electron microscope, were larger, but also showed that they were significantly less numerous. Staining of neuronal nuclei revealed that this reduction in synapse number was not due to a reduction in the number or density of neurons, since depending on the measurement, these were either unchanged in number or were modestly increased (**Supplemental Figure 2**). Because synapses between excitatory neurons in neocortex are thought to occur primarily or exclusively at dendritic spines, we also measured linear spine density as a second light microscopic method of assaying changes in excitatory synaptic density. These measurements revealed a significant reduction in the number of synapses per unit dendritic length, providing indirect corroboration for the observation of reduced synapse density.

Prior studies have conflicted on the issue of whether or not blocking synaptic transmission causes spine loss. Some studies have suggested that blocking glutamate receptors leads to spine and synapse loss, while blocking action potentials does not (McKinney et al., 1999), while other studies have observed a loss of synapses following chronic (Mitra et al., 2011), but not acute (24 h) activity blockade with TTX (Kuhlmann et al., 2021) and still others have found synapse loss after both chronic (Hazan and Ziv, 2020) and acute (Minerbi et al., 2009) blockade of activity. Even studies manipulating release genetically come to differing conclusions about the quantitative impact, with one study of Munc 13 double knockout slice cultures concluding that spine density was unchanged despite complete loss of spontaneous and evoked transmission (Sigler et al., 2017) while another study performed on slice cultures in which tetanus toxin expression led to a similar loss of vesicular release concluded that although many spines formed normally, there were ^40% fewer spines and synapses in CA1 (Sando et al., 2017). Differences in preparations studied, or in the manipulations used may account for some of the differences in results. However, in studies reporting synapse loss, this is typically accompanied by an increase in the strength of the remaining synapses (Hazan and Ziv, 2020; Minerbi et al., 2009; Mitra et al., 2011).

Although not directly measured here, we know that after five days of deprivation there is a strong activity rebound reflecting circuit hyperexcitability. Our experiments suggest that one factor contributing to this hyperexcitability is likely enhanced excitatory synaptic transmission, as evidenced by increased frequency and amplitude of mEPSCs and enlarged pre- and postsynaptic size. The fact that the functional increase in frequency is seen despite a reduction in apparent synapse number suggests a strong increase in presynaptic release probability at the remaining synapses, consistent with the observation of greater numbers of docked vesicles and with measures of release probability undertaken after activity blockade in other systems (Han and Stevens, 2009). Although we focused only on excitatory synapses in this study, it is likely that the enhanced network excitability is augmented by increased intrinsic excitability (Desai et al., 1999) and by decreases in inhibitory synaptic transmission (Kilman et al., 2002).

One of our motivations for performing measurements of the size of synapses at the ultrastructural level and the size of puncta of synaptic proteins in the light microscope was the hope of validating the latter as an approach for assaying structural properties of synapses in dense tissue with relatively high throughput. We developed and validated automated quantitative methods that may prove useful for future studies of synapse size and number. Our approach shares a number of features with that used in a study published recently (Sauerbeck et al., 2020). This approach, also made use of Airyscan super resolution imaging of synaptic proteins, and employed thresholding and image processing via the Imaris imaging platform, and demonstrated a reduction in synapse number similar in magnitude to that seen in separate samples using electron microscopy. Our approach could potentially be further improved with additional image processing, such as the 2D sliding window thresholding, template matching, and background subtraction, used by the PunctaSpecks algorithm (Shah et al., 2020), or with machine learning approaches as in the DoGNet system (Kulikov et al., 2019). Although the lower resolution of even super-resolution light microscopy makes it poorly suited to identify structural changes at individual synapses, it may nevertheless be suited for studying the effects of more global manipulations on the distributions of synaptic properties as done here.

## SUPPLEMENTAL FIGURES

**Supplemental Figure 1.**
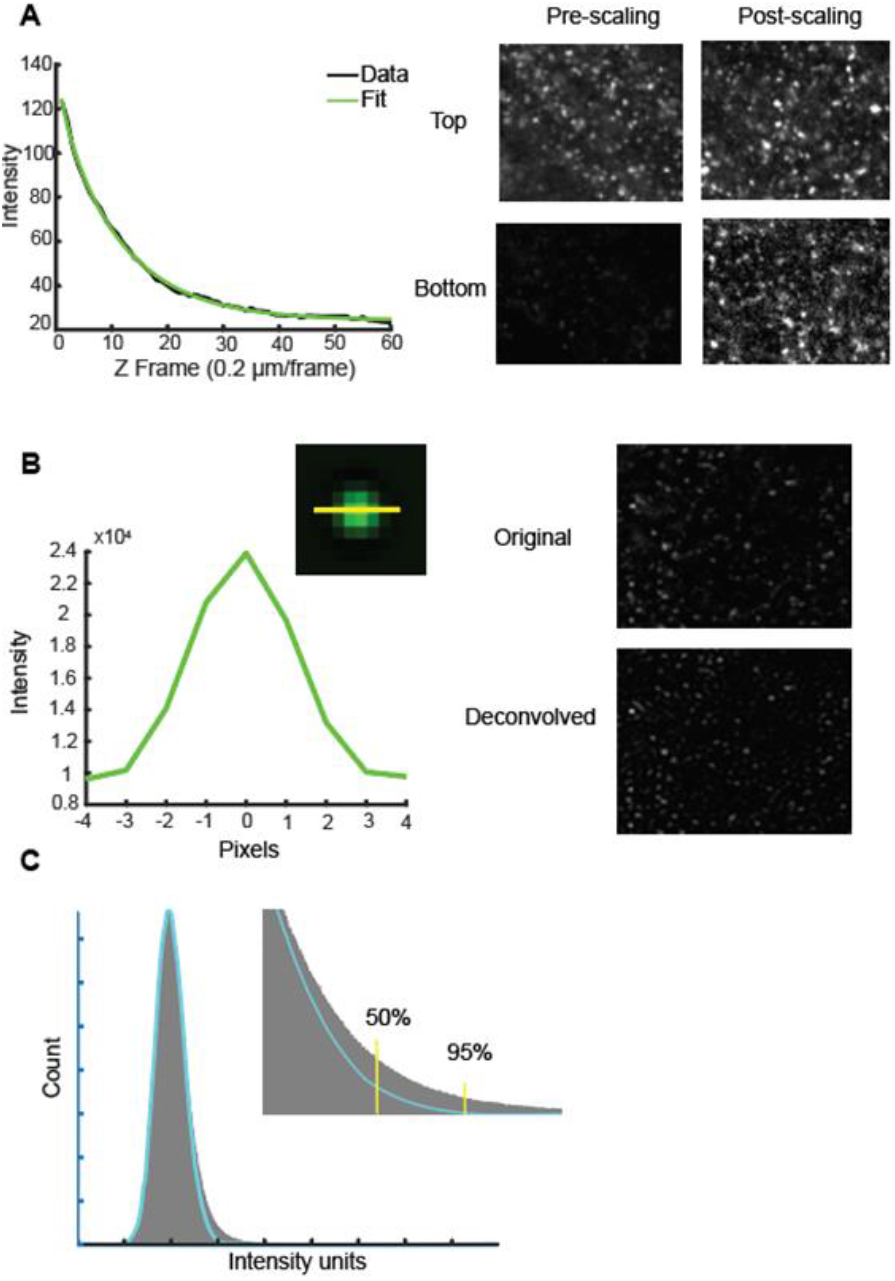
Methods for analysis of light microscopy data from Airyscan. **A**. Example of light scattering in tissue. Left shows an example before scaling where, over 60 μm, signal is drastically reduced to the point where a contrast appropriate for the first frame gives essentially a black field by the last frame. The scaling fit over depth (green) of this data (black) is shown in the middle graph. The same stack rescaled is shown on the far right. **B**. Left: A 100 nm diameter fluorescent bead in green, with 1D line drawn through the center, yields a point-spread function graph. Right: Example transformation from deconvolution of a single channel example image using 10 such beads averaged into a kernel. **C**. Thresholding by signal-to-noise ratio. Histogram of all intensity values in dark gray, with skewed normal distribution fit to the maxima of this histogram in cyan. Inset shows two potential signal-to-noise percentage based thresholds for illustration in yellow, 50% (where 50% of the values here are likely to be part of noise, and 50% are likely to be signal) and 95% (5% chance noise, 95% chance signal).

**Supplemental Figure 2.**
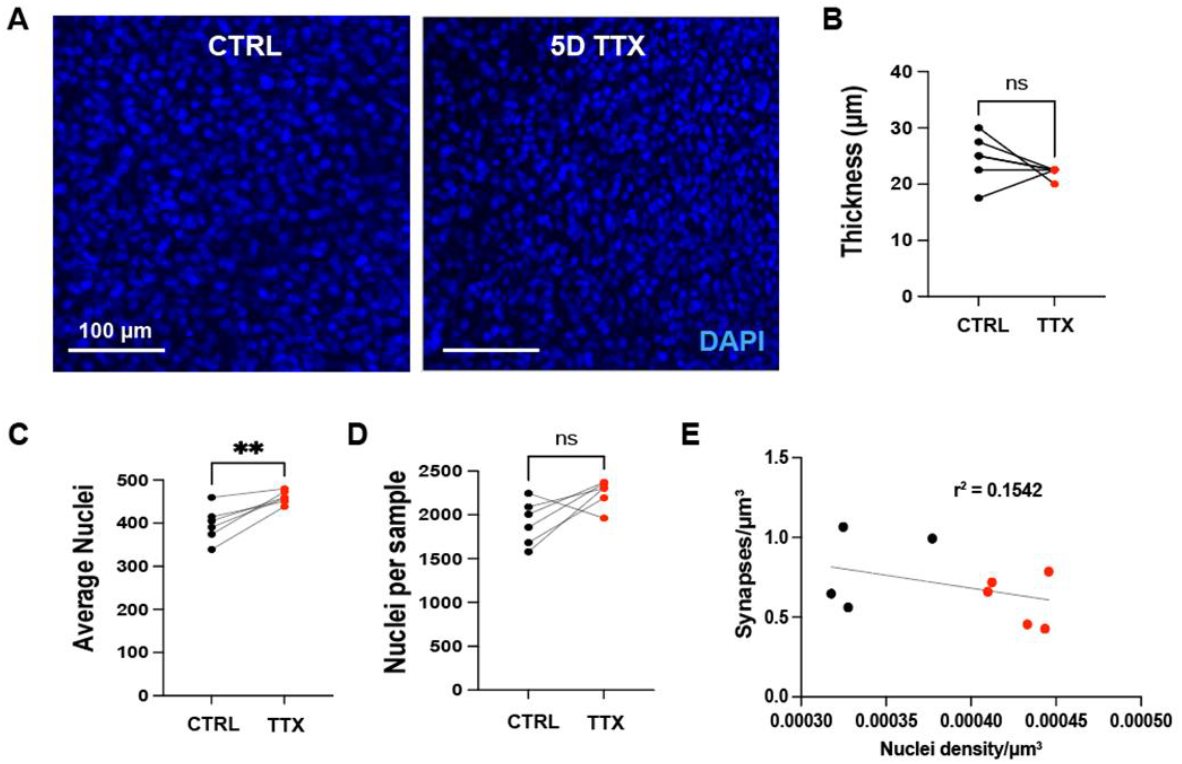
Prolonged silencing does not decrease cell density. **A**. Example 10X confocal image from untreated and TTX-treated slices. Scale bar = 100 μm. **B**. Quantification of slice thickness of control and TTX-treated samples. **C**. Average DAPI stained nuclei found per 2.5 μm z-frame (same region as **Figure 4F**). **D**. Sum of DAPI nuclei in the full thickness (22-25μm) imaged in these samples. **E**. Relationship between synapse density and nuclei density for samples from 4F; CTRL (red) and TTX-treated (black).

